# Spatial mapping of posture-dependent resistance to passive displacement of the hypertonic arm post-stroke

**DOI:** 10.1101/2022.11.13.516311

**Authors:** Priyanka Kanade-Mehta, Maria Bengtson, Tina Stoeckmann, John McGuire, Claude Ghez, Robert A Scheidt

## Abstract

Muscles in the post-stroke arm commonly demonstrate abnormal reflexes that result in increased position- and velocity-dependent resistance to movement. We sought to develop a reliable way to quantify mechanical consequences of abnormal neuromuscular mechanisms throughout the reachable workspace in the hemiparetic arm post-stroke. Survivors of hemiparetic stroke (HS) and neurologically-intact (NI) control subjects were instructed to relax as a robotic device repositioned the hand of their (hemiparetic) arm between several testing locations sampling the arm’s passive range of motion. During transitions, the robot induced motions at either the shoulder or elbow joint at speeds ranging from very slow (6 °/s) to fast (90 °/s). The robot held the hand at the testing location for at least 20 seconds after each transition. We recorded and analyzed hand force and electromyographic activities from selected muscles spanning the shoulder and elbow during and after transitions. Hand forces and electromyographic activities were small at all speeds and sample times in NI control subjects, but varied systematically by transport speed during and shortly after movement in the HS subjects. Velocity-dependent resistance to stretch diminished within two seconds after movement ceased in the hemiparetic arms. Hand forces and EMGs changed very little from 2 seconds after the movement ended onward, exhibiting dependence on limb posture but no systematic dependence on movement speed or direction. Although each HS subject displayed a unique field of hand forces and EMG responses across the workspace after movement ceased, the magnitude of steady-state hand forces was generally greater near the outer boundaries of the workspace than in the center of the workspace for the HS group but not the NI group. In the HS group, electromyographic activities exhibited abnormalities consistent with stroke-related decreases in the stretch reflex thresholds post-stroke. These observations were consistent across repeated testing days. Implications of the findings are discussed.

## Introduction

Muscles in the post-stroke arm commonly demonstrate abnormal reflexes that result in increased position- and velocity-dependent resistance to movement (i.e. spastic hypertonus: Lance, 1980; Li et al., 2021). Spastic hypertonia is understood to reflect systematic reductions in stretch reflex thresholds (Katz and Rymer, 1989; Levin et al., 2000), decreased range of regulation of these stretch reflex thresholds (Levin and Feldman 1994), as well as altered non-reflex phenomena such as abnormalities in the intrinsic mechanical properties of spastic muscles and altered viscoelastic properties of passive tissues (Mirbagheri et al., 2007; Sinkjaer and Magnussen 1994; Zhang et al., 2002). Importantly, systematic reduction in stretch reflex threshold could lead to significant increase in stretch reflex excitability (Knutsson and Owens 2003) and agonist/antagonist coactivity in some regions of the workspace (Levin and Dimov, 1997; Musampa et al., 2007), which could lead to complex, posture-dependent and potentially time-varying joint impedances in the hemiparetic arm. Because the feedforward control of goal-directed movements relies on accurate predictions of limb impedance (Scheidt et al., 2011), spatial and temporal complexity of joint impedance may be a significant contributor to impairment of movement coordination post-stroke (Scheidt and Stoeckmann, 2007; Zackowski et al. 2004). Although a velocity-dependent increase in muscle tone (spasticity) is frequently assessed clinically and has been quantified in single joints such as the elbow and wrist (Levin et al.1994; Li et al. 2005; Katz and Rymer; 1992; Starsky et al, 2005) assessments of multijoint control deficits have been rare (see Musampa et al. 2007; Sangani et al. 2007). The goal of this study was to develop a reliable and repeatable approach for measuring the mechanical consequences of abnormal neuromuscular mechanisms as a function of hand location in the reachable workspace in the hemiparetic arm post-stroke. We designed a set of experiments using a two-joint, planar robot to quantitatively measure the mechanical and electromyographic responses to controlled displacements of the upper extremity at several locations in the arm’s workspace. Subjects were instructed to relax as the robot moved their hand sequentially between target locations spanning the reachable workspace. Trajectories were selected such that movements were largely limited to either the shoulder or elbow joint, but not both. Three different joint transport velocities were programmed, ranging from very slow (6°/s) to fast (90°/s). The robot stabilized the hand at the target for at least 20 seconds following the end of each movement. We analyzed the time series of horizontal planar hand forces and electromyographic (EMG) activities for selected arm muscles to assess the contribution of abnormal reflex and non-reflex mechanisms to phasic and tonic (steady-state) components of posture-dependent joint torques post-stroke. Specifically, we wished to determine the duration of phasic, velocity-dependent resistance to stretch, to characterize the spatial topography of tonic, position-dependent hand forces throughout the reachable workspace, and to characterize the muscle activities that give rise to these postural bias forces. The resulting data demonstrate that the robotic assessment of posture-dependent bias forces was repeatable across days, that the phasic component of these stroke-related forces lasted no more than two seconds after the end of limb re-positioning, and that these bias forces were partly neuromuscular in origin (not merely due to passive tissue resistance to stretch) such that elevated “resting” EMG activity exhibited posture-dependence in some muscles, but posture-invariance in others. Ramifications of the findings are discussed.

## Methods

A convenience sample of ten unilateral, hemiparetic survivors of stroke (HS; aged 43-62 years) and eleven neurologically intact control subjects (NI; age-range matched: 43-61 years) gave informed consent to participate in this study (Table 1). All procedures were approved by Marquette University’s Office of Research Compliance in accord with the Declaration of Helsinki. All HS were recruited from the pool of hemiparetic stroke outpatients of Medical College of Wisconsin and the Milwaukee VA Medical Center and all were in the chronic stage of recovery (between 2 and 28 years post-stroke). Exclusion criteria for HS included: inability to give informed consent, inability to follow 2-step directions, history of tendon transfer in the affected limb, physical dimensions prohibiting appropriate interaction with the robotic assessment system (Fig 1) despite reasonable adjustment attempts (e.g. abdominal intrusion into the robot’s workspace or an inability to see the feedback display after chair height adjustment), use of aminoglycoside antibiotics, curare-like agents, or other agents that might interfere with neuromuscular transmission, botulinum toxin treatment within the previous 8 months, and/or shoulder pain in the test position of 75° to 90° abduction. The presence of contracture or shoulder subluxation did not exclude subjects from participating unless it limited their ability to perform the experiments comfortably. NI control subjects were required to have no history of neurological disorder. All subjects were able to achieve all test positions without discomfort. All NI subjects were right-handed. Three of the NI subjects participated in three experimental sessions with at least one week separating each session. The remaining eight NI subjects participated in a single session. Seven HS participated in two experimental sessions separated in time by at least one week. The remaining HS participated in three experimental sessions on separate days. Each session lasted ∼2 hours. Handedness was assessed for all subjects using the Edinburgh Handedness Inventory (EHI). HS were asked to use pre-stroke preferences to guide their answers. Table 1 provides a description of subject characteristics for all subjects. Within the group of survivors, FM_UE_ scores ranged from 19 to 53, reflecting a broad spectrum of functional motor impairment within our subject pool.

**Table 1:** Subject characteristics

| Subject | Sex | Age | Year post stroke | Stroke Type | Paretic Side | MAS | Impairment | FM | Elbow |  | Shoulder |  | Handed |
| --- | --- | --- | --- | --- | --- | --- | --- | --- | --- | --- | --- | --- | --- |
|  |  |  |  |  |  |  |  |  | min flex | max ext | min flex | max ext |  |
| HS01 | F | 54 | 9 | I | L | 1.25 | Mild | 53 | 60 | 175 | 70 | 160 | R |
| HS02 | F | 58 | 25 | H | L | 2 | Severe | 19 | 60 | 175 | 70 | 160 | R |
| HS03 | F | 63 | 5.7 | I | L | 0.5 | Mild | 40 | 60 | 175 | 70 | 160 | A |
| HS04 | F | 53 | 5.9 | H | L | 2.5 | Moderate | 27 | 60 | 175 | 70 | 160 | R |
| HS05 | M | 49 | 6 | I | L | 2 | Severe | 23 | 35 | 95 | 80 | 130 | L |
| HS06 | M | 59 | 2.10 | I | R | 1 | Moderate | 38 | 100 | 40 | 100 | 30 | R |
| HS07 | M | 52 | 28 | H | L | 1.75 | Severe | 24 | 80 | 165 | 70 | 160 | R |
| HS08 | F | 53 | 12 | I | L | 2.5 | Moderate | 31 | 80 | 165 | 70 | 160 | R |
| HS09 | M | 61 | 3 | I | R | 3.5 | Moderate | 29 | 100 | 40 | 100 | 30 | R |
| HS10 | M | 64 | 8 | I | L | 2 | Mild | 43 | 80 | 165 | 70 | 160 | R |
| NI01 | M | 43 |  |  |  |  |  |  | 100 | 40 | 100 | 30 | R |
| NI02 | F | 52 |  |  |  |  |  |  | 100 | 40 | 100 | 30 | R |
| NI03 | M | 49 |  |  |  |  |  |  | 100 | 40 | 100 | 30 | R |
| NI04 | M | 45 |  |  |  |  |  |  | 100 | 40 | 100 | 30 | R |
| NI05 | M | 63 |  |  |  |  |  |  | 100 | 40 | 100 | 30 | R |
| NI06 | F | 60 |  |  |  |  |  |  | 100 | 40 | 100 | 30 | R |
| NI07 | F | 58 |  |  |  |  |  |  | 100 | 40 | 100 | 30 | R |
| NI08 | M | 58 |  |  |  |  |  |  | 100 | 40 | 100 | 30 | R |
| NI09 | F | 55 |  |  |  |  |  |  | 100 | 40 | 100 | 30 | R |

**Stroke Type:** I: Ischemic stroke, H: Hemorrhagic Stroke; **MAS:** Modified Ashworth Score, averaged across joints [e.g. if elbow flexion = 1 and elbow extension = 1+, then MAS = (1+1.5)/2]; **FM:** Fugl Meyer Score, upper extremity portion (max = 66); **EF:** Elbow Flexion; **EE:** Elbow Extension; **SF:** Shoulder Flexion; **SE:** Shoulder Extension; **Handedness:** R: right; L: left; A: ambidextrous.

**Figure 1:**
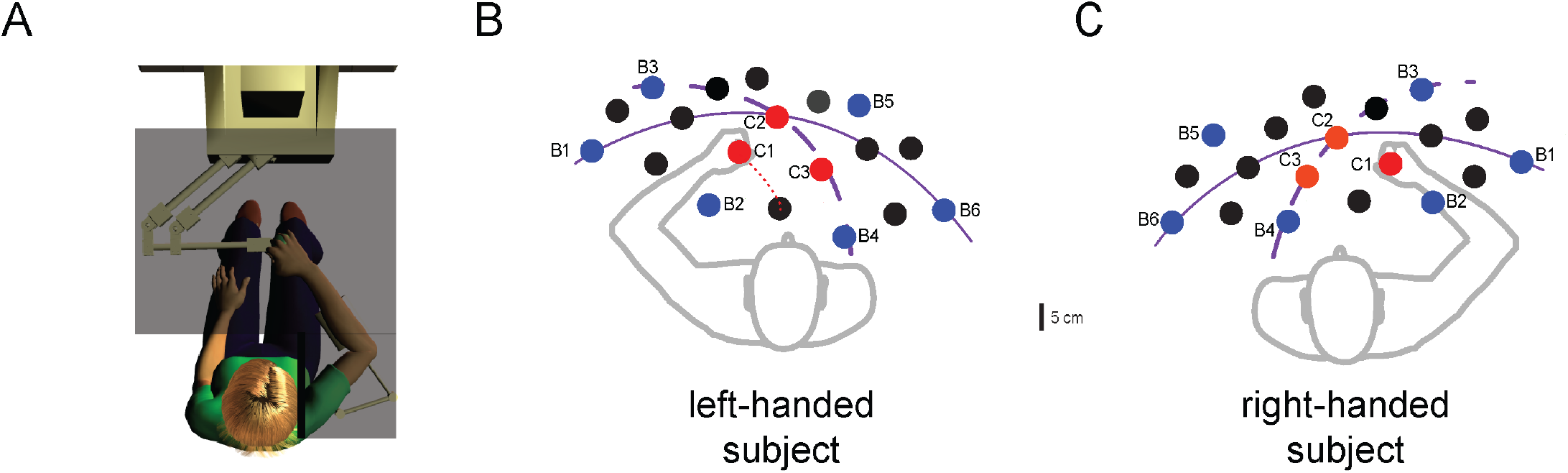
Experimental setup. **A**) Subject seated at the horizontal planar robotic testing system. The system included the robotic tool as well as a horizontal planar feedback display and electromyography sensors (not shown). **B** and **C**) Representative left-hand and right-hand workspaces with sample points. Selected sample locations are labeled as boundary (B1 through B6, blue circles) or center points (C1 through C3, red circles). Interior points are shown in black. The left- and right-hand workspaces were mirror-symmetric such that the hand was at boundary point B4 when the shoulder and elbow both were flexed to the maximum extent allowed by the subject’s passive range of motion and/or the robot’s workspace.

### Clinical assessments

Prior to experimentation, HS participated in a series of clinical assessments. The upper extremity portion of the Fugl-Meyer assessment of motor performance (FM_UE_) was used to assess impairment in the ability of each HS to move the more affected arm (Fugl-Meyer 1975). Scoring for the FM_UE_ is on a 0-66 point scale, which is based the subject’s ability to move both within and outside of muscle synergies. Increasing values reflect greater motor control, especially of the distal limb segments. The Modified Ashworth Scale (MAS) was used to assess hypertonia in the flexors and extensors at the shoulder, elbow, and wrist. Scoring for the MAS is on a 0-4-point scale with 0 indicating no difference between the more- and less-affected sides and a 4 indicating that the more-affected limb is rigid. Within each limb, MAS scores were averaged across the three joints tested to obtain an overall estimate of upper extremity spasticity (range: 0–4; see also Zackowski et al. 2004). Passive range of motion of the elbow and shoulder in the workspace plane was measured using a goniometer both with and without the robotic test apparatus.

### Experimental procedures

Subjects were seated in an adjustable, high-backed chair fixed in front of an actuated, 2-joint robotic manipulandum designed to move in a horizontal plane (see Scheidt et al., 2010 for details). Subjects were secured into the chair using a seatbelt-style chest harness that served to minimize trunk motion. Efforts were taken to situate subjects in the same location and orientation relative to the robot’s workspace from one testing day to the next. An opaque screen mounted 1 cm above the plane of hand motion occluded vision of the arm, hand, and robotic manipulandum. The subject’s wrist was splinted at 0° flexion and fixed at the wrist to the robot’s handle with Velcro straps. The arm was supported against gravity (between 75° and 90° abduction angle) using a lightweight, chair-mounted arm support. Arm segment lengths and passive range of motion at the shoulder and elbow were used to determine each individual’s reachable workspace in the horizontal plane. A minimum of 18 spatial sample locations (targets) spanned the full range of shoulder and elbow flexion and extension (Fig 1B and 1C) except in subjects with reachable workspaces exceeding the robot’s workspace. In those cases, the sample locations spanned the intersection of the subject’s range of motion and that of the planar robot. For each subject, detailed analyses were performed at three targets defined as center locations (red dots labeled C1-C3 in Figs 1B and 1C) and at six targets defined as boundary locations (blue dots labeled B1-B6 in Figs 1B and 1C). The remaining points were used to construct spatial maps of posture-dependent interface forces and muscle activities as described below. Subjects were verbally coached throughout the experiment to “relax” at all times and not to intervene as the robot generated passive movements of the arm through its workspace. To reinforce our verbal coaching, the word RELAX was projected onto the occluding screen throughout the experimental session.

The robot monitored instantaneous hand position using rotational encoders (A25SB17P180C06E1CN; Gurley Instruments Inc., Troy, NY) mounted on the motor shafts (M-605-A Goldline; Kollmorgen, Inc. Northampton, MA). The location of the handle could be resolved within 0.038 mm throughout the operating workspace. The robot was programmed to generate stiff control of hand position at a rate of 1000 samples/sec. A 16-bit data acquisition board (PCI-6031E DAQ; National Instruments Inc., Austin, TX) sampled analog force data from a load cell (85M35A-I40-A-200N12; JR3 Inc., Woodland, CA) that was mounted immediately under the handle.

The robot enforced transitions between targets such that motion occurred predominantly at one joint (the focal joint: either the shoulder or elbow) while the non-focal joint remained essentially still. Figs 1B and 1C show elbow and shoulder trajectories (dotted and solid arcs, respectively) along which only the focal joint was flexed or extended.

Hand trajectories had bell-shaped speed profiles computed such that rotation of the focal joint had peak angular velocities that were slow (6°/s), medium (30°/s), or fast (90°/s). In contrast to step- or ramp-shaped velocity profiles, a bell-shaped profile achieves smoother transitions between one location and another, imitating the smooth changes in arm position applied during manual testing of muscle resistance by clinicians (Levin et al., 2000). After the end of transition (EOT), the robot held the hand at the target location for at least 20 seconds (the “holding period”). Targets C1-C3 were visited from all four directions (elbow extension, elbow flexion, shoulder extension, and shoulder flexion) in pseudorandom order. Targets located at a workspace boundary (B1-B6) were visited from at least two directions in pseudorandom order. In total, 105 transitions were used to visit each target from all desired directions at the different speeds (also pseudorandomized across trials).

### Data Collection and Analysis

We recorded instantaneous hand position and force as well as surface electromyograms (EMG) at a rate of 1000 samples/s. Monitored muscles included: shoulder horizontal adductors pectoralis major (PECS) and anterior deltoid (ADL), shoulder horizontal abductor posterior deltoid (PDL), elbow flexors biceps (short and long heads: BICS and BICL) and brachioradialis (BRD), as well as elbow extensors triceps (lateral and long heads: TRILT and TRILG). Raw EMGs were amplified x1000 (Myosystem 1200, Noraxon, Inc. Scottsdale, AZ) and band-pass filtered between 10 and 500 Hz in hardware prior to digitization. 60 Hz and 120 Hz artifacts were eliminated in post-processing using zero-phase, 4^th^-order Butterworth notch filters. Residual offsets were removed from the digitized EMGs prior to rectification and low-pass filtering at 4 Hz with a zero-phase, 4^th^-order Butterworth filter.

To facilitate comparison of EMG activity across the study population, each subject’s muscle activities were normalized by the peak value of the rectified and filtered activity recorded from that muscle during a series of 5 s duration maximum voluntary isometric contractions (MVICs). MVICs were performed with the hand stabilized in the center of the arm’s horizontal planar workspace. Each subject performed 12 MVIC trials before the experiment: three each of maximal isometric elbow flexion, elbow extension, shoulder horizontal abduction and adduction. The peak magnitude of EMG activity for each muscle was defined as the largest EMG value for that muscle within any of the MVIC trials after signal processing as described in the preceding paragraph. We also recorded 5 s of quiet resting activity in the same limb configuration prior to MVIC recording and after approximately 5 minutes of rest, wherein subjects were encouraged to “relax”.

A primary goal of our analysis was to differentiate velocity-dependent resistance to stretch post-stroke from position-dependent, steady-state effects. To this end, we filtered the kinematic and kinetic time series off-line using a zero lag, 4^th^ order, 10 Hz Butterworth low-pass filter and analyzed the hand’s force vector at nine points in time including the moment of peak hand speed (V_MAX_), the moment of target acquisition (End of Transition or EOT, defined as the time when hand velocity first dropped below 15% of its peak value), as well as at 1, 2, 3, 5, 10, 15, and 20 seconds after EOT. We also analyzed the magnitude of rectified and filtered EMG signals at the same time points.

### Statistical testing

We performed two main analyses. The first analyzed interface forces at the robot’s handle to quantify velocity-dependent resistance to passive motion of the arm. We controlled for posture-dependencies so as to determine the earliest point in time after EOT beyond which hand force ceases to demonstrate velocity-dependence. This was done to better isolate posture-dependent effects in the second analysis described below. As we will show, hand forces measured at the end of the holding period displayed no systematic dependence on movement speed, and so we regarded hand forces measured at EOT+20 s as representative of asymptotic values. We subtracted these values from hand forces measured at other time points in the same trial before performing a six-way, repeated measures, general linear model ANOVA examining how changes in measured hand forces (relative to EOT+20 s) varied by subject type (HS or NI), movement speed (6⯚/s, 30⯚/s, 90⯚/s), sampling instant (time of peak transit velocity, EOT, and 1, 2, 3, 5, and 15 seconds after EOT), movement direction (elbow flexion, elbow extension, shoulder flexion, shoulder extension), workspace location (center vs. boundary, wherein the center locations were collapsed across C1-C3 and the boundary locations were collapsed across B1-B6) and testing day (1-3). Post-hoc ANOVA and Tukey t-tests were performed to examine significant main and interaction effects.

The second analysis characterized the spatial topography of tonic, position-dependent hand forces as a function of workspace location at the earliest time point wherein hand forces neither depended on transport velocity nor differed from the asymptotic values. To do so, we plotted raw hand force vectors as a function of workspace location for each subject. These plots were then co-registered with contemporaneous maps of normalized EMG activities to determine whether the observed hand forces might be partly neuromuscular in origin. Specifically, we evaluated the extent to which shoulder and elbow muscle activities varied as a function of joint angle along the solid (PECS, ADL, and PDL) and dashed lines (BICS, BICL, BRD, TRILT, and TRILG) in Figs 1B and 1C.

Data processing and statistical testing were carried out within the Matlab (Mathworks, Inc., Natick, MA) and Minitab (Minitab Inc., State College, PA) computing environments. Effects were considered statistically significant at the α =0.05 level. Specifically, we applied a Bonferroni correction to the 20 statistical tests performed on the hand force data, which yielded an individual test significance threshold of p = 0.0025. Despite consistency in the mechanical recordings, the EMG data demonstrated considerable variation across stroke survivors. Consequently, we did not correct the EMG analyses for multiple comparisons (i.e. we accepted statistical significance at p = 0.05).

## Results

All subjects were alert throughout each experimental session even though they were not required to engage in any task-dependent activity aside from relaxing. Robot-generated hand movements were always smooth, having unimodal, bell-shaped velocity profiles (Fig 2A). Passive displacement of the hand modulated EMG activity in HS subjects (especially in stretched muscles) but rarely did so in NI control subjects. For example, stretching shoulder or elbow muscles in the hemiparetic arm caused EMG activity in some muscles to increase during movement (Fig 2, left), as might be expected due to spastic hypertonia (i.e. Schmit and Rymer, 2001). Interestingly, elevated muscle activity could remain active long after motion had ceased in some regions of the workspace, as might be expected due to a decrease in the stretch reflex threshold post-stroke (Levin and Feldman, 1994), although other explanations are possible. In some muscles, tonic EMG activity after passive displacement of the hand appeared to depend on the final limb configuration (posture). Persistence of muscle activity was never observed in NI control subjects during or after passive movement (cf. Fig 2, right).

**Figure 2:**
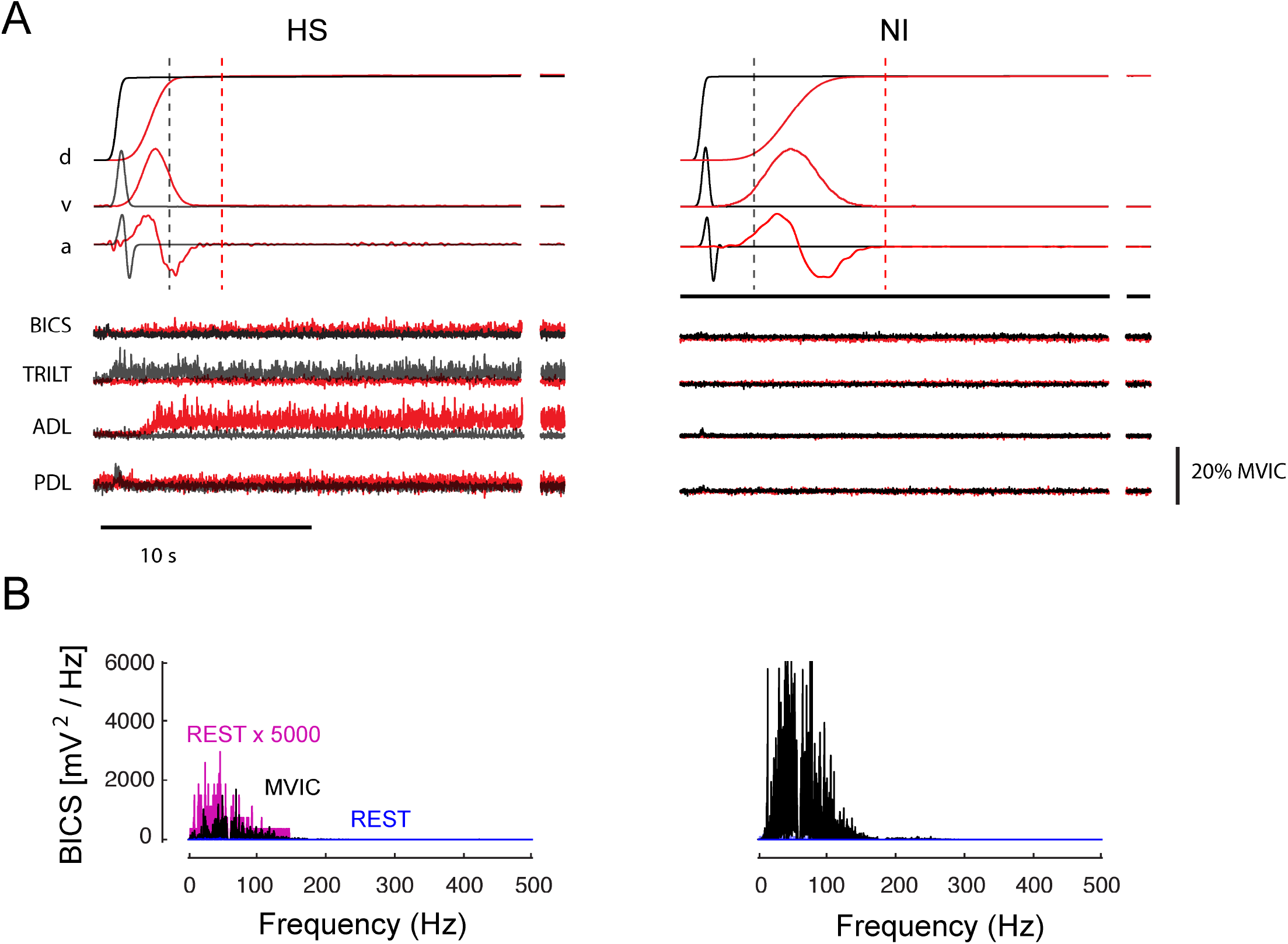
A) Representative normalized movement kinematics (top) and normalized, rectified EMG signals (bottom) as a function of time for a single fast transition between targets C1 and B4 (grey) and a single moderate speed transition between targets C2 and B1 (red) in a representative SS subject (left). A single fast transition between targets C1 and B4 (grey) and a slow transition between targets C2 and B1 (red) in a representative NI control subject is shown on the right. Horizontal scale bar: 10 s. Vertical scale bar: EMG signal amplitude of 20% MVIC. Movement kinematics are displayed on an arbitrary scale to highlight general characteristics such as smoothness. The additional snippets of data to the right of each trace present data from the very end of the hold period (at least 25 s after the end of movement). Vertical dashed lines indicate the time of EOT + 2 s. B) Spectrograms of BICS EMG signal power during MVIC trials (black) and quiet rest (blue) for the same subjects. Also shown for comparison for the HS subject (left) is the resting spectrogram multiplied by a factor of 5000 (purple).

Elevated EMG activities in HS subjects were not an artifact due to poor signal transduction or improper EMG normalization in these subjects. Because EMG normalization with respect to MVIC could overestimate voluntary muscular effort if the signals were corrupted by environmental noise, we visually verified that EMGs recorded from each muscle were of high quality by plotting the EMG signal power spectrograms from MVIC trials for each muscle and compared it to spectrograms obtained during quiet rest. In all cases, MVIC EMG signal power was distributed across the frequency range in a unimodal pattern characteristic of high-quality surface EMG recordings (Roman-Liu and Konarska 2009) (Fig 2B, black). By contrast, EMG signal power was lower during rest in the center of the workspace (Fig 2B, blue). MVIC EMG exceeded quiet resting signal power by a factor greater than 100 in all cases (>> 20 dB signal to noise ratio). Thus, normalization with respect to MVIC yielded a high-quality assessment of relative voluntary effort. Note that while resting EMG power was uniformly negligible at all frequencies in NI subjects, it commonly displayed measurable, tonic activity post-stroke with a spectral signature of quality surface EMGs rather than broadband environmental noise (e.g. Fig 2B, left; purple).

We sought to characterize how mechanical resistance to passive joint motion varied with movement velocity (i.e., spastic hypertonia; Lance 1980). Visualization of raw hand forces from a selected HS subject (Fig 3A) demonstrated that hand force magnitude could vary systematically by transport speed during movement, that variations in steady-state hand force were not systematic during the hold phase that followed, and that hand forces changed very little from EOT + 20 s onward. Despite efforts to minimize trunk motion, some trial-by-trial variability in hand force undoubtedly arose from subjects shifting in their seat during the ∼1.5-hour testing period. Because the effect of these infrequent postural shifts would be to alter the bias forces recorded at the handle throughout the trial, we reasoned that greater sensitivity in subsequent analyses of velocity-dependence would be achieved by removing the effects of postural shifts (i.e. by analyzing hand forces relative to their asymptotic values). Indeed, exploratory three-way repeated measures ANOVA found that hand force magnitude at EOT+20 s post-stroke did not vary systematically with transport speed (F_(3,207)_=1.40, p=0.25). We therefore aligned the raw hand force profiles in time with respect to EOT and subtracted asymptotic values on a trial-by-trial basis (Fig 3B) prior to evaluating how subject group, workspace location and other factors influence the transient effect of movement on reaction forces at the hand.

**Figure 3:**
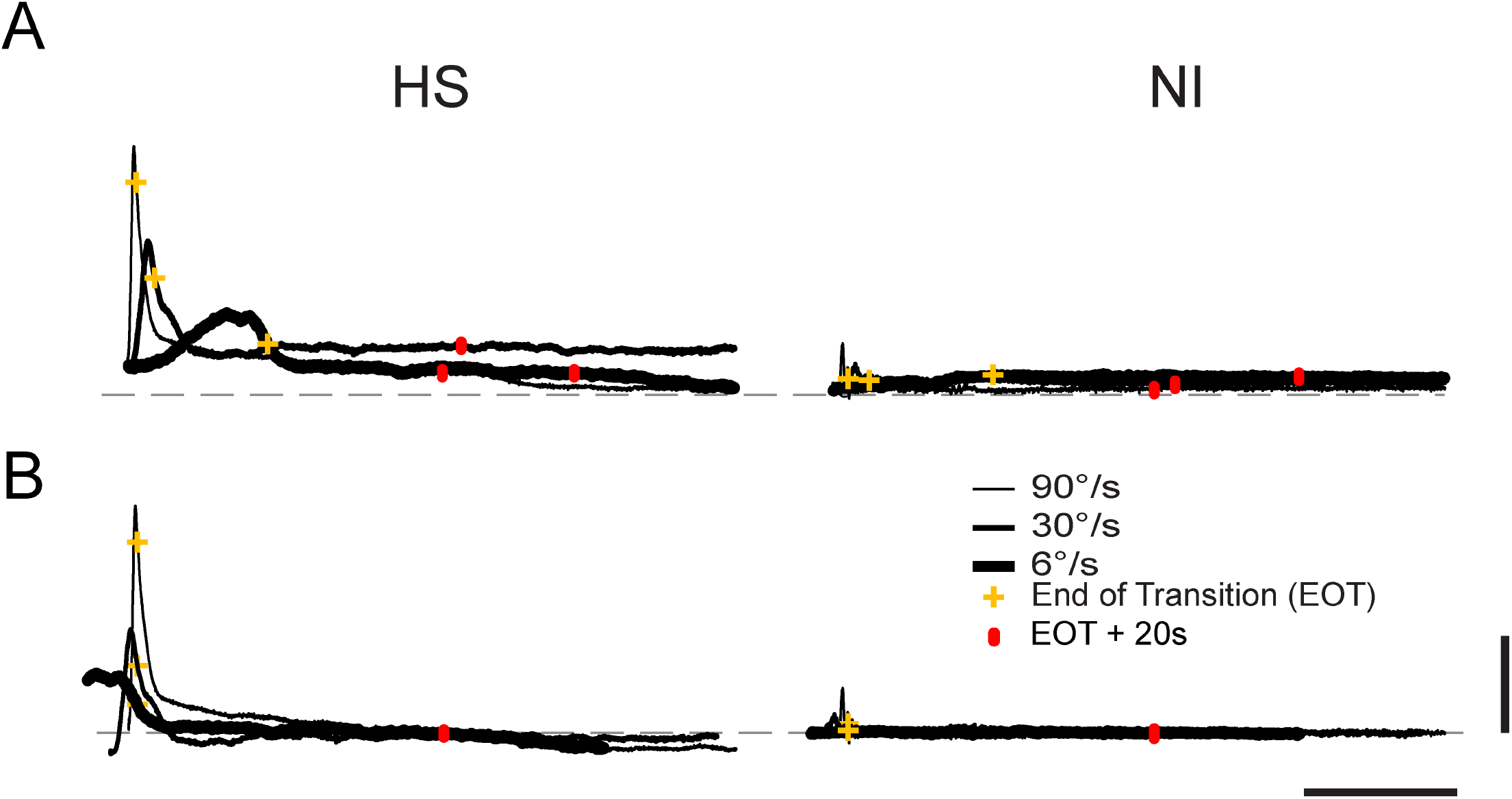
A) Hand forces as a function of time during transitions between targets as indicated by the red trajectory in Figure 1 (panel B) for a representative SS subject (left) and NI control subject (right) at three transport speeds (90⯚: black; 30⯚: blue; 6⯚: green). Markers (yellow +) indicate the time of End Of Transition (EOT). Markers (red hashes) indicate EOT + 20s. Vertical scale bar corresponds to a measured hand force of 20N while the horizontal dashed line indicates 0 measured hand force. The horizontal scale corresponds to 10 s. B) Force profiles plotted with respect to steady-state values (EOT+20s) and aligned in time with respect to EOT. Here, the vertical scale bar corresponds to a change in measured hand force of 20N *relative to steady-state*.

### Hand force measurements are repeatable across days

We performed a seven-way, mixed-model, general linear model, repeated measures ANOVA to quantify how changes in measured hand force magnitude (relative to EOT+20 s) varied by subject group, movement speed, movement direction, workspace location (boundary vs center), temporal sampling instant and testing day in response to passive relocation of the subject’s hand from one workspace location to another. We found that testing day failed to demonstrate a significant main effect (F_(2, 4799)_ = 0.26, p = 0.769) and that this factor also had no interaction with any other factor (p > 0.21 in each case). Moreover, 78% of the Day 2 variations in hand force (across subjects, workspace location, speed, movement direction and sampling window) were predicted by measurements made on Day 1. From this we conclude that measurement of endpoint forces using our experimental approach was repeatable across days.

### Effect of movement speed on measured hand forces

Although the ANOVA found no main effect of movement direction or workspace location on hand force (relative to steady-state) and no significant interaction between subject group and either of these factors, the analysis did find a significant three-way interaction between subject group, sampling instant and movement speed (F_(14, 4799)_ = 3.02; p<0.0005). No other three-way or higher-order interactions achieved significance, indicating that the observed velocity-dependent effects did not vary systematically across the workspace after stroke. To test whether the observed three-way interaction could possibly have been due to alterations in the passive viscoelastic properties of tissues spanning the shoulder and elbow joints post-stroke, we repeated the ANOVA on the sampled hand force data from the 30⯚/s and 90⯚/s trials after subtracting values obtained during the 6⯚/s trials (i.e. trials wherein velocity-dependent stretch reflex activity – but not viscoelastic resistance - should have been minimal; Schmit and Rymer 2001; Starsky et al. 2005). We obtained similar results and identical statistical conclusions from this supplemental analysis (results not shown). Thus, velocity-dependent responses measured during and shortly after movement were not solely due to passive tissue viscoelasticity but rather implicated the presence of abnormal stretch reflexes post-stroke.

Figure 4 plots change in hand force magnitude relative to steady-state asymptote as averaged across target locations, movement directions and days. The three-way interaction between subject group, sampling window and movement speed can readily be seen in that hand forces differed across groups, transport speeds, and sampling times during and shortly after transport but not later in the holding period. To determine the earliest point in time beyond which hand force ceased to demonstrate movement velocity-dependence, we performed a post-hoc series of eight separate two-way, mixed-model, general linear model, repeated measures ANOVA to determine how changes in hand force magnitude varied by subject group and movement speed for each of the eight sampling times (PeakVel, EOT, and EOT plus 1, 2, 3, 5, 10 and 15 seconds). In contrast to the first three sampling times, wherein the main effect of movement speed was either significant (PeakVel: F _(2,38)_ = 70.58, p< 0.0005; EOT: F_(2,38)_ = 21.14, p < 0.0005) or marginally significant (EOT+1: F_(2,38)_ = 2.76, p = 0.085), the main effect of movement speed was absent at EOT+2s (F_(2,38)_ = 0.91, p = 0.417) and at all subsequent time points. As shown by the vertical dashed lines in Fig 2A, arm movement had completely ceased prior to EOT+2s (i.e. there was no acceleration or velocity component associated with EOT+2s for any movement speed). The interaction between movement speed and subject group was significant only during movement (PeakVel: F _(2,38)_ = 8.44, p< 0.002); no interactions were found at later times. A set of one-sided t-tests revealed that hand force magnitude post-stroke did not differ from asymptotic values from EOT+2s onwards. Thus, spastic responses (i.e. velocity-dependent resistance to stretch) are most visible in the post-stroke hand in the fastest conditions of limb transport, but only *during* transport or for a short period thereafter (i.e. less than 2 seconds after movement ceases).

**Figure 4:**
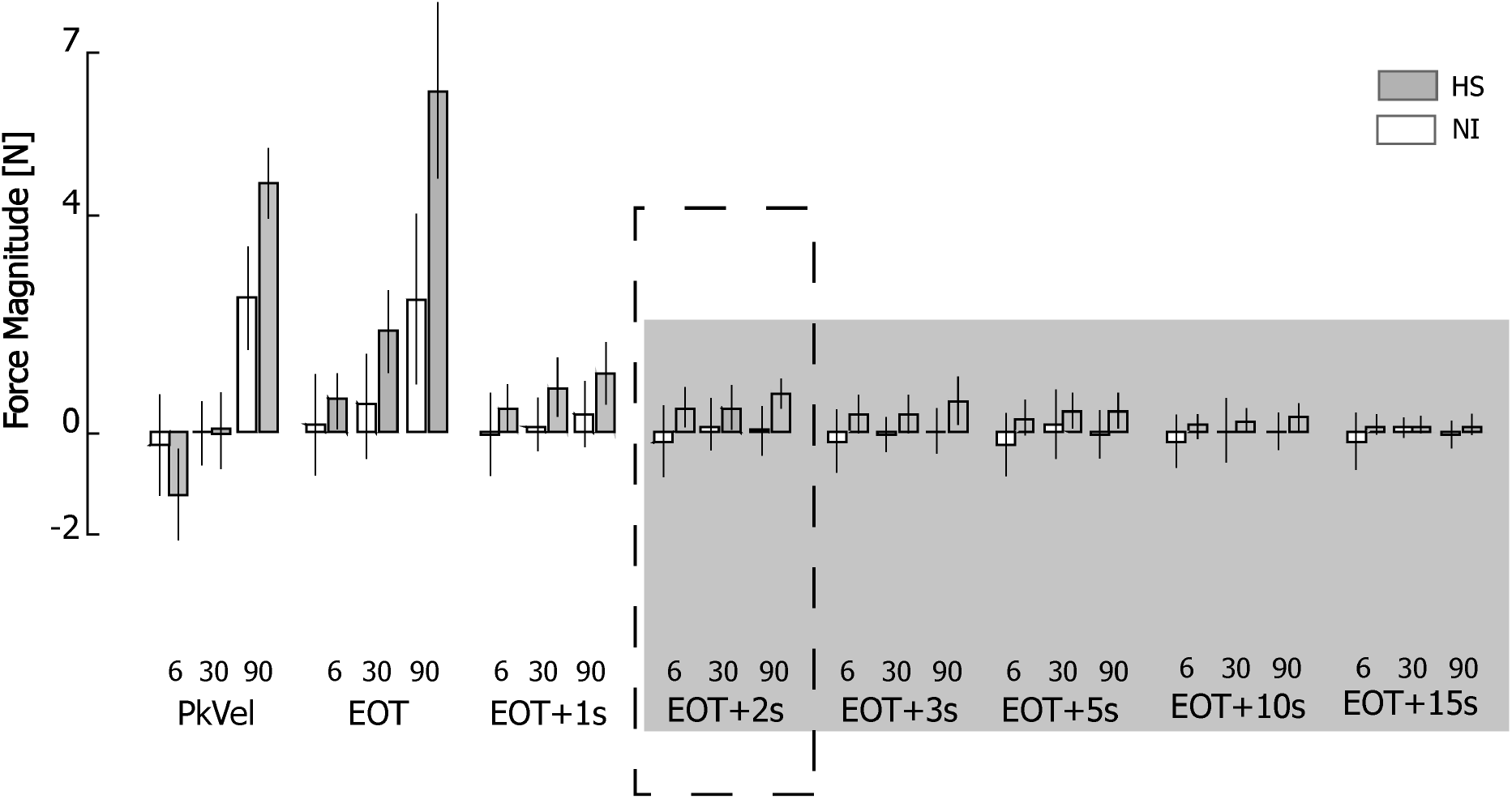
Force magnitude as a function of speed, time and subject group (HS: Shaded bars; NI: open bars). Error bars represent ±1 SEM. The shaded area from EOT+2 s onwards indicates the sampling intervals in which ANOVA found no effect of movement speed on hand force magnitude.

### Effect of hand position on postural bias forces

Because hand forces reached velocity-independence and steady-state by EOT+2s, we next plotted the raw hand forces - not relative to asymptote - measured at EOT+2s for each subject. Representative HS and NI subject data are shown in Fig 5A. Compared to NI subjects who generated negligible tonic forces at the robot’s handle throughout the workspace after movement had ceased, each HS subject displayed a unique field of hand forces that varied systematically across their arm’s reachable workspace. Steady-state, posture-dependent hand forces were typically higher at the workspace boundaries in HS subjects. These forces were generally stronger on one side of the reachable workspace and pointed towards an equilibrium point located in the approximate center of the workspace.

**Figure 5:**
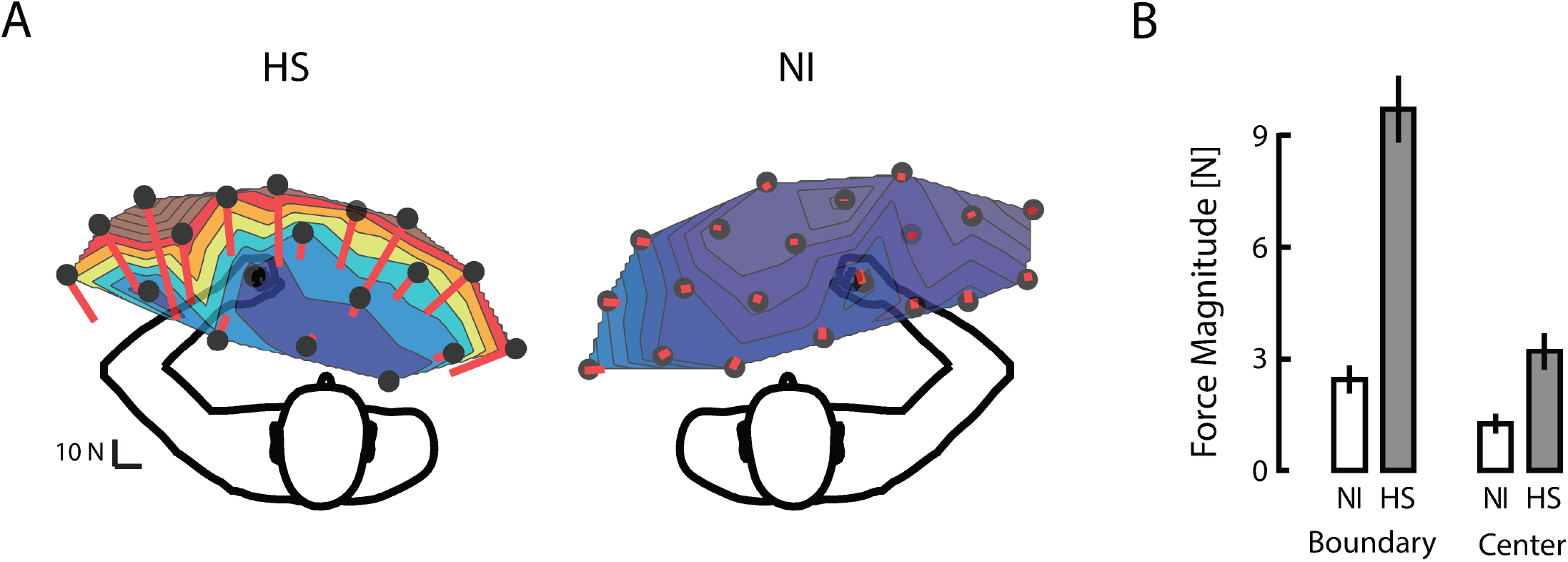
A) Raw hand force vectors as a function of location throughout the workspace for a representative HS subject (left) and NI subject (right). B) Population summary of hand force magnitude as a function of workspace location (boundary vs. center) for HS (grey bars) and NI subjects (open bars) at EOT+2s. Error bars represent ±1 SEM.

We used a mixed-model, repeated measures ANOVA to examine how hand force magnitude values measured at EOT+2 s varied as a function of subject group (HS, NI), movement direction (EF, EE, SF, SE), and workspace location (boundary, center). The dependent variable (raw hand force magnitude; not relative to asymptote) was averaged across movement speeds and days. Workspace location demonstrated a strong interaction with subject group (F_(1, 37)_ = 51.24, p<0.0005): hand forces varied strongly depending on whether the location was at or near the boundary or center of the workspace for HS (Fig 5B) but not NI subjects, for whom statistical significance did not survive Bonferroni correction for multiple comparisons. The ANOVA found no evidence supporting a main effect of movement direction or any interaction between this and the other factors (p > 0.301 in all cases). Thus, hand forces measured at EOT+2 s post-stroke were predominantly position-dependent, having no systematic dependence on movement speed and/or direction.

### Elevated hand forces are accompanied by large EMG activity in some hemiparetic arm muscles post-stroke

We next sought to determine whether posture-dependent bias forces post-stroke were due primarily to passive properties of tissues spanning the hemiparetic joints or whether bias forces were at least partly neuromuscular in origin. Fig 6 plots spatial maps of selected EMG activities (normalized to MVIC) at EOT+2 s for representative subjects from each group. For the selected HS subject, robotic translation of the hand led to elevated levels of EMG activities that were relatively large in some muscles with respect to signals recorded during maximal voluntary isometric contractions (e.g., TRILT, TRILG and BRD). In other muscles, the activities tended to exhibit posture-dependence such that activity was greater when the muscle was lengthened than when shortened (e.g., BICL, PECS). Yet other muscles exhibited negligible activities at EOT+2s regardless of limb posture (e.g., BICS, ADL, PDL). These results were characteristic of the study population in the sense that all stroke survivors exhibited high levels of muscle activity throughout the workspace only in some muscles (most notably TRI and PECS), a modest tendency to exhibit posture-dependent activity in a one or two muscles (which varied by individual), and no indication of abnormal “resting” activity in the remaining muscles. By contrast, activity was minimal in all muscles at EOT+2s throughout the workspace for all NI subjects (a selected individual’s results are shown in Fig 6B).

**Figure 6:**
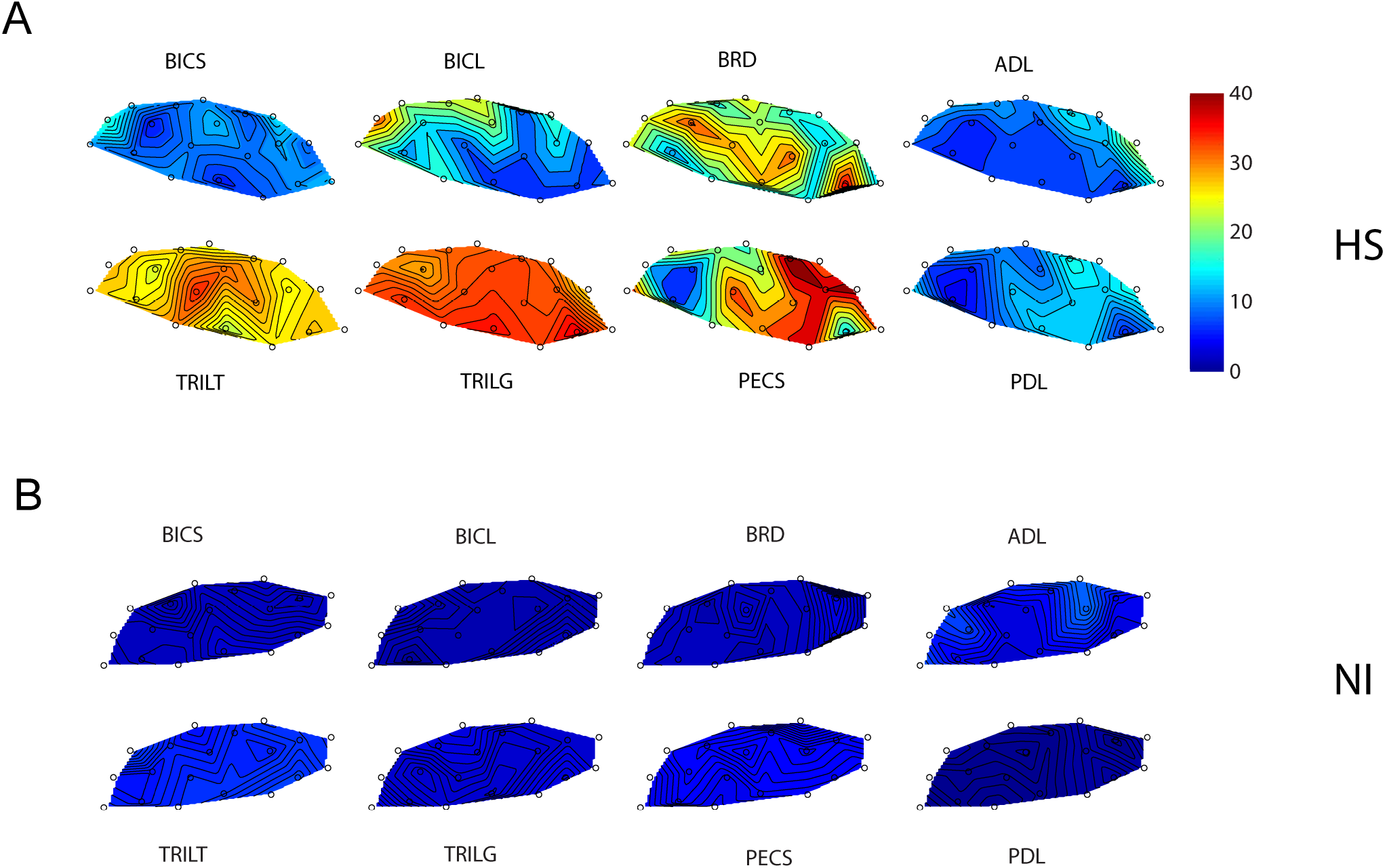
A) Contour plots of elbow and shoulder muscle activities (averaged across transition speeds) for a selected HS subject at 2 s after End of Transition. Muscle activation is presented as a function of hand position in the workspace on a scale ranging from 0 to 100% maximum isometric voluntary contraction (color bar on the right). B) Contour plots of elbow and shoulder muscles for a selected NI subject at 2 s after End of Transition.

We performed separate sets of three-way, repeated measures, general linear model ANOVA to examine the extent to which elbow and shoulder muscle activities varied by subject group {NI, HS}, sample time {EOT+2 s, EOT+20 s}, and workspace location. We focused the analyses on positions along the dashed arc (elbow analyses) and solid arc (shoulder analyses) shown in Figs 1B and 1C. For the elbow analyses (Fig 7, top), we found no main effect related to sample time (F_(1,115)_ < 0.68; p > 0.412 in all cases), or any interaction between sample time and the other two fixed factors (F_(1,115)_ < 0.93; p > 0.336 in all cases). Consistent with our hypothesis that interaction forces induced by passive translation of the hand are partly neuromuscular in origin, we found a main effect of subject group for BRD (F_(1,115)_ = 8.67; p = 0.010), TRILG (F_(1,115)_ = 12.66; p = 0.003), and BICL (F_(1,115)_ = 6.68; p = 0.020). In each case, the measured muscle activities were a larger percentage of their voluntary maximum capacity throughout the workspace for the HS group as compared to the NI control group. We did not observe a main effect of subject group for BICS or TRILT (F_(1,115)_ < 0.71 and p > 0.411 in both cases). We did however observe an apparent interaction between subject group and target for BRD (F_(3,115)_ = 3.86; p = 0.011) such that for the HS group only, BRD activity was systematically greater when the elbow was extended and that muscle was stretched, vs. when the elbow was flexed and the muscle was shortened. We observed no other two-way interactions for any of the recorded elbow muscles.

**Figure 7:**
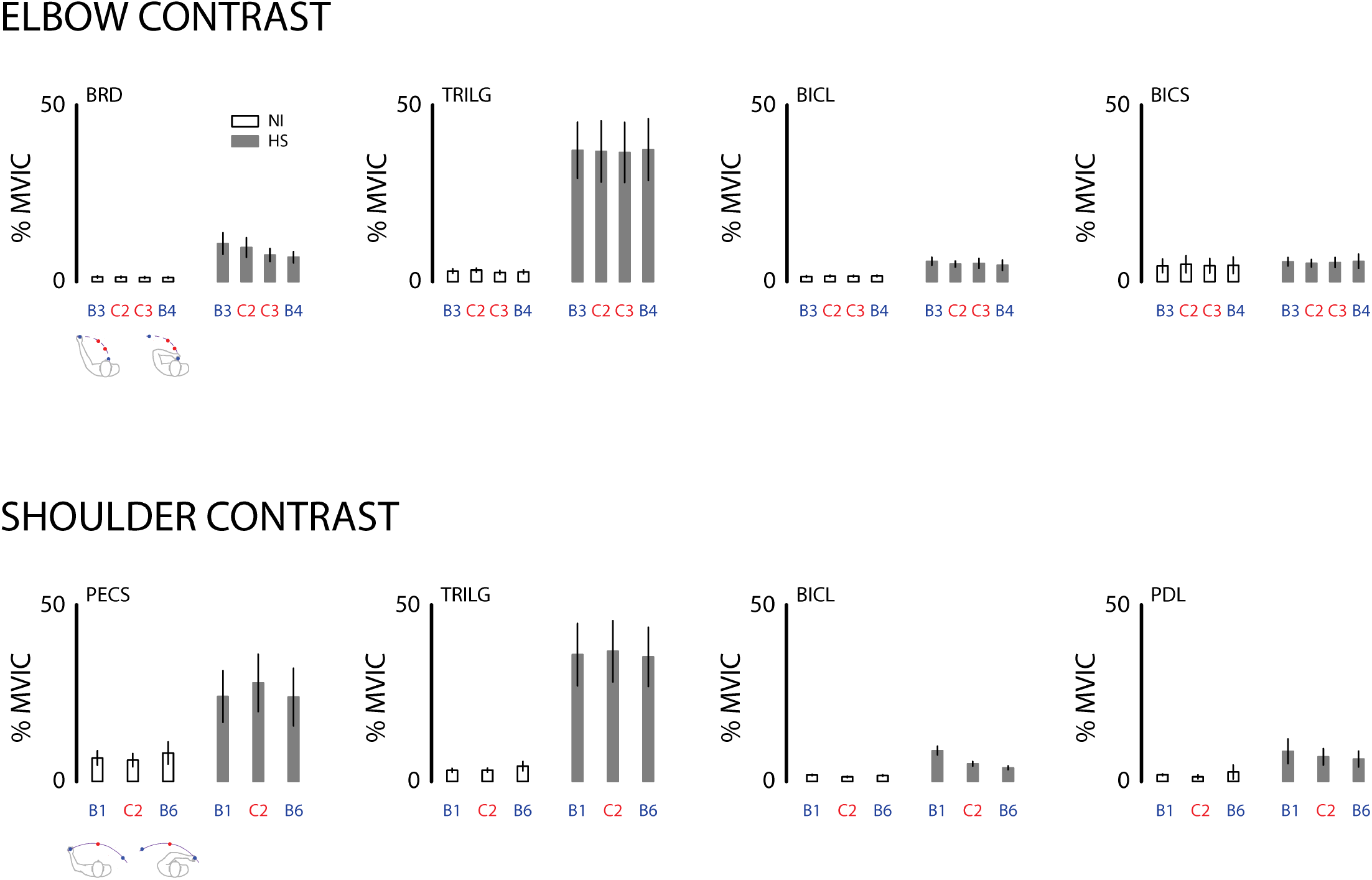
Cohort results: analyses of selected muscle activities (as a percentage of MVIC) at EOT+2 s in the elbow contrast (top) and shoulder contrast (bottom), as described in the text. B: workspace boundary positions; C: central positions. Icons below the left panels depict limb configurations for the two boundary targets in each contrast. Grey bars: HS; Open bars: NI. Error bars represent ±1 SEM.

Likewise for the shoulder analyses (Fig 7, bottom), we found no main effect related to sample time (F_(1,82)_ < 0.97; p > 0.328 in all cases), or any interaction between sample time and the other two fixed factors (F_(1,82)_ < 0.93; p > 0.338 in all cases. We found a main effect of subject group for PECS (F_(1,82)_ = 4.70; p = 0.046, TRILG (F_(1,82)_ = 11.12; p = 0.004), and BICL (F_(1,82)_ = 10.98; p = 0.004). Here again, the measured muscle activities were a greater percentage of MVIC throughout the workspace for the HS group when compared to the NI group. We did not observe a systematic main effect of subject group for PDL or ADL (F_(1,82)_ < 3.01 and p > 0.102 in both cases). We observed an interaction between subject group and target for BICL (F_(2,82)_ = 24.24; p < 0.0005) such that for the HS group only, BICL activity was greater when the shoulder was extended and that muscle was stretched, vs. when the shoulder was flexed and the muscle was shortened. We observed no other two-way interactions for any of the recorded shoulder muscles.

We examined further the two significant interaction effects by calculating for each subject an *EMG modulation index* as the difference between normalized EOT+2 s EMG values measured at the target location where the muscles were most flexed vs. where the muscles were most extended (Fig 8). For BRD (elbow contrast), this entailed subtracting normalized EMG values measure at target B4 from those measured at target B3. For BICL (shoulder contrast), this entailed subtracting values measure at target B6 from those measured at target B1. EMG modulation index values were tightly packed around 0% MVIC in the NI group. By contrast, index values were greater than zero in both the BRD and BICL muscles in the HS group. While a single HS appeared to be an outlier in both cases (BRD: HS09; BICL: HS02) (Fig 8; open squares), removing these outliers and repeating the analyses described above did not impact the pattern of main and interaction effects reported above. Thus, we obtained support for the idea that passively stretched muscles held in an elongated vs. shortened state elicit greater involuntary activations after stroke in a subset of tested muscles. Neither modulation index exhibited significant correlation with either FMUE or MAS scores (p>0.09 in all four cases).

**Figure 8:**
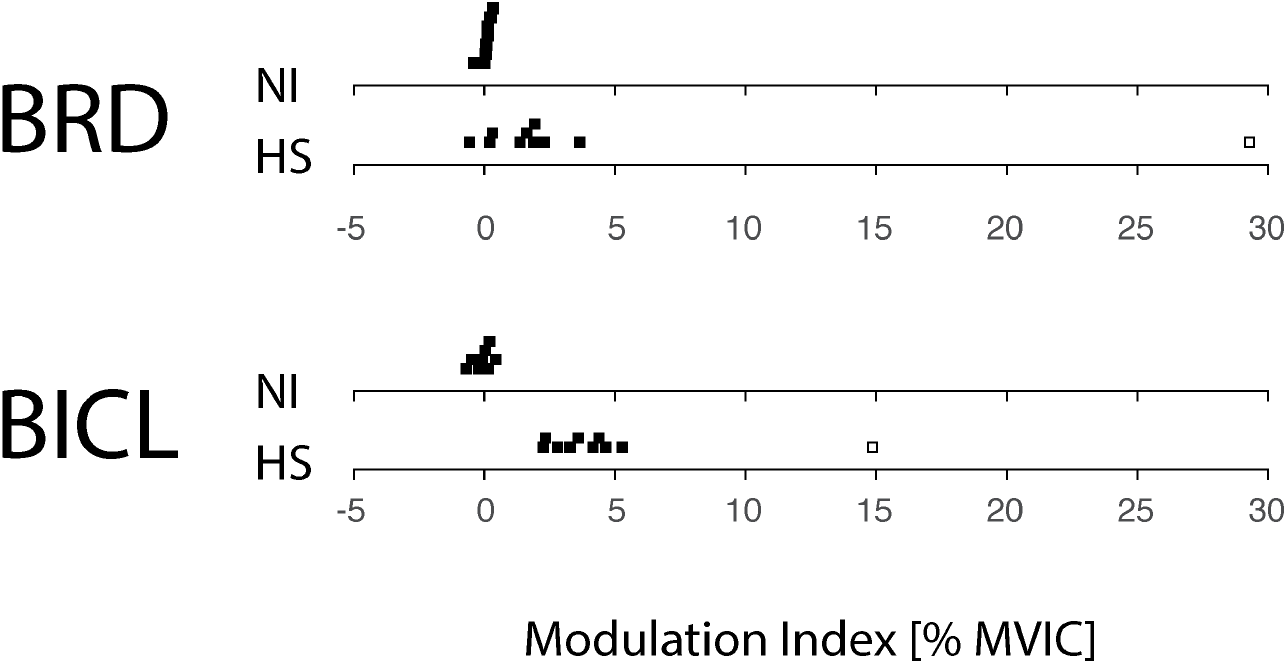
Cohort results: EMG Modulation Index [i.e., the change in normalized EMG values measured across the workspace for the elbow contrast (BRD) and the shoulder contrast (BICL) as described in the text]. Each dot represents the modulation index value obtained from a single subject in either the group of NI control subjects or the group of HS subjects.

## Discussion

We used a planar robot to assess the spatial and temporal topography of position-dependent hand forces and electromyographic activities that arise during and after passive movement of the arm after stroke. Survivors of hemiparetic stroke (HS) and neurologically intact control subjects (NI) were instructed to relax as the robot repositioned the hand at several different testing locations within a horizontal planar slice through the passive range of motion for the (hemiparetic) arm. The robot was programmed such that transitions between testing locations induced motion primarily at either the shoulder or elbow joint (but not both) at three different speeds ranging from very slow (6°/s) to fast (90°/s). The robot held the hand at the testing location for a minimum of 20 seconds after each transition. We recorded hand force and electromyographic activities in selected muscles spanning the shoulder and elbow joints throughout the transition and during the holding period that followed. All of the HS and a subset of the NI subjects returned to the lab on subsequent days to repeat the testing procedures one to two times. We found that hand interaction forces were relatively small at all speeds and sample times in NI subjects, whereas they varied systematically by transport speed during and shortly after movement in HS subjects. Observation of velocity-dependence after movement had ceased indicates that hand forces were not solely due to passive tissue viscoelasticity post-stroke, but rather implicated the presence of abnormal stretch reflexes. In the HS group, spastic responses (i.e. velocity-dependent resistance to stretch) diminished within two seconds after the end of transition (EOT). Hand forces changed very little from EOT+2s to the end of the sampling period (EOT+20s), exhibiting dependence on limb posture but no systematic dependence on movement speed and/or direction. Although each HS subject displayed a unique field of hand forces across the arm’s reachable workspace after movement had ceased, the magnitude of steady-state hand force was generally greater near the outer boundaries of the workspace than in the center of the workspace for the HS group but not the NI group. In the HS group, hand forces were generally stronger on one side of the reachable workspace and pointed towards an equilibrium posture located in the approximate center of the workspace. These observations were consistent (repeatable) across testing days. We consider the steady-state hand forces to reflect abnormal postural biases because they depend on history of stroke and limb configuration but not on the speed and direction of passive movements bringing the hand to the different spatial sampling locations.

Stroke-related postural bias forces were partly neuromuscular in origin because they were typically accompanied by persistently elevated EMG activity levels in most muscles spanning the shoulder and elbow joints. During and shortly after movement, passive translation of the hemiparetic arm led to increased EMG activities during movement consistent with spastic hypertonia (e.g., Schmit and Rymer, 2001). In some muscles, elevated muscle activity could remain active long after motion had ceased, as might be expected due to a decrease in the stretch reflex threshold post-stroke (Levin and Feldman, 1994; Levin et al 2000). These neuromuscular responses to passive displacement of the hand were persistent in the sense that the complex spatial patterns of EMG activities observed at EOT+2 s were also observed at EOT+20s. The specific details as to which muscles exhibited increased activities varied from person to person, although some general trends were apparent. Across subjects in the HS group, muscles that exhibited posture dependent activities at EOT+2 s also did so at EOT+20 (BRD, BICL). Muscles that exhibited elevated activity throughout the workspace at EOT+2s also did so at EOT+20s (TRILG, PECS). Muscles that did not exhibit elevated activities at EOT+2 s also failed to do so at EOT+20 s (BICS, TRILT, ADL, PDL). By contrast, persistence of muscle activity was never observed in NI control subjects during or after passive movement (cf. Fig 2, right; Fig 7).

### Disentangling velocity- and posture-dependent responses to imposed arm movement - transient and steady state effects

Several prior studies have examined transient mechanical and electromyographic responses to passive displacements of the wrist or elbow to quantitatively assess post-stroke spasticity (cf., Katz et al 1992; Levin and Feldman 1994; Li et al 2006; Starsky et al 2005). In one example, Schmit and colleagues used a motorized device to passively flex and extend the elbow of hemiparetic stroke survivors over a range of speeds ranging from slow (6º/sec) to fast (90º/sec) (Starsky et al., 2005). Their goal was to assess the reliability of three different biomechanical correlates of spasticity, which they isolated from other aspects of spastic hypertonia associated with dystonia, contracture, and increased joint stiffness. They did so by subtracting the torque response to the slowest displacement from responses to faster displacements, leaving only reflex torque. This approach is effective because the stiffness of the passive tissues about the elbow joint are largely velocity insensitive (Given et al. 1995). Of the three biomechanical measures considered - peak torque, peak joint stiffness, and onset angle of reflex torque responses - peak torque values measured during displacements at 90°/sec were most reliable on repeated measures in a single testing session (>80% reliability), and most highly correlated with clinical assessments of spasticity (Ashworth Scale). In another example, Mirbhageri and colleagues (Mirbhageri et al. 2008) used system identification technique to quantify the contributions of reflex and intrinsic (i.e., non-reflex) stiffness to total elbow stiffness at several different elbow angles in the paretic and nonparetic arm of chronic hemiparetic stroke survivors. Each position was examined under passive conditions in the range of full elbow flexion to full elbow extension. They reported that intrinsic and reflex stiffness both contributed strongly to net joint torque (equivalent to endpoint hand forces observed in our study), that the effects were significantly larger in the paretic than in the non-paretic elbow muscles, and that these differences increased with the increasing joint angle indicating position dependence. Schmit and Rymer (2001) created a muscle activation model to measure the static and dynamic components of the stretch reflex observed during passive extension of the elbow at 10 different velocities by examining different aspects of the torque/angle relationship. They found that four out of six model parameters reflected the static stretch reflex response and the remaining two reflected the dynamic stretch reflex response. The mechanism of static tonic stretch reflexes presumably involves receptors that are chiefly sensitive to muscle length and not velocity. The secondary muscle spindle afferents maintain an increased firing over baseline for as long as the muscle is held stretched and would be suitable candidates. Muscle spindle primaries are suitable candidates for mediating the dynamic effects.

Our findings extend the results of these previous studies in that hand forces in steady state after displacement were almost exclusively position dependent, with no measurable contributions attributable to movement speed, direction or repetition of across days. Though we also observed significant velocity-dependent responses during and shortly after passive translation of the hand, the current study focused on steady-state mechanical and electromyographic responses rather than on transient responses because prior studies of goal-directed reaching found that deficits in the control of limb posture and movement were dissociable after stroke (Scheidt and Stoeckmann 2007; Mani et al. 2013). We therefore sought to characterize the spatial topography of position-dependent hand forces and electromyographic activities that arise even with the arm at rest. We observed no residual velocity-dependence in hand forces measured at time points 2 seconds after the end on hand translation and later. We also observed no systematic differences between hand forces measured at EOT+2s and EOT+20s. We therefore analyzed the steady-state reaction forces measured at the hand at EOT+2s and found systematic increases in posture-dependent hand force in some regions of the workspace but not others for all subjects in the HS group. Although the fine details of the spatial topography of hand forces were subject-dependent, we did observe systematic variations such that hand reaction forces were greater along the boundary of the arm’s workspace than in the center of the workspace (Fig 5B). These effects were repeatable in the sense that similar results were obtained with repeated testing on separate days. The increased bias forces were neuromuscular in origin - at least in part - because the regions of elevated hand forces corresponded with regions of elevated muscle activity in some of the muscles spanning the shoulder and elbow joints (e.g., Fig 7; BRD and BICL; Fig 8). Although other muscles also exhibit abnormal activations throughout the workspace (e.g., TRILG, PECS, PDL), the measured bias forces reflect a mechanical contribution from elbow flexors (as in Fig 5A) consistent with the expression of the classic abnormal flexion synergy (Dewald et al. 1995).

These findings raise several questions that are ripe for further study. For example, the range of muscle activations available for voluntary control should be greater in regions of lower bias force than in regions where bias forces are greater. If so, then there should be systematic interactions between the bias forces measured with the limb at rest and the ability to perform voluntary actions throughout the arm’s workspace. Is it easier to stabilize the limb against environmental perturbation in regions of low bias force rather than in regions where bias forces are greater in magnitude? Might moving into a region of greater bias force be more difficult (and less accurate) than moving into a region of lower bias force? Preliminary data from a pilot study suggest that such interactions may indeed arise after stroke (Simo et al., 2013; Laczko et al., 2017).

### Spatial topography of postural biases

Others have also studied the spatial topography of EMG activities elicited by passive displacements of the arm post-stroke, albeit with limited spatial resolution. Using “quasi-static” (<5º/s) passive stretches applied to the elbow starting from full elbow flexion or extension and with the shoulder held at three joint angles, Musampa and colleagues (2007) sought to determine the elbow joint angles at which the different flexor and extensor muscles would begin to be activated. They observed the static stretch reflex thresholds encroached on the active range of motion post-stroke and varied significantly with initial shoulder angles. This effect was not observed in neurologically intact control subjects. The authors attributed these observations to the presence of a tonic stretch reflex response at rest due to elevated excitability of α-motor neurons innervating stretched muscles arising from a loss of descending inhibition and the ensuing imbalance of excitation and inhibition (cf., Lance 1980; Powers et al. 1988). A consequence of the encroachment of static stretch reflex thresholds into the arm’s typical range of motion is the existence of spasticity zones - i.e., regions of the arm’s workspace in which some flexor or extensor muscles cannot be relaxed. For some muscles, spasticity zones occupied a substantial part of the biomechanically defined range of motion. In some cases, the spasticity zones of antagonistic flexor–extensor muscle groups overlapped, yielding what the authors called “spatial co-activation zone” (Musampa et al., 2007). Other studies have reported that abnormal muscle coactivations and joint torque-coupling patterns constrain the ability of individuals with stroke to voluntarily generate the full typical range of joint torque combinations (Ellis et al. 2007; see also Sangani et al. 2007). Abnormal patterns of agonist and antagonist muscle activation during stretch or voluntary movement have been previously described in single- and double-joint muscles around the elbow joint in adults and children with hemiparesis (Dewald et al., 1995; Beer et al., 2000; Dewald and Beer, 2001; Levin et al., 2000; Jobin and Levin 2000; Sangani et al. 2007).

The results of our present study confirm and extend the findings of those prior studies, demonstrating that position-dependent hand forces and EMGs observed in the static hold phase post stroke in our study are indeed neuromuscular in origin - at least in part. Across our cohort of HS subjects (Fig 7), we observed all three patterns of stretch reflex threshold variations predicted by Musampa et al. (2007). In some muscles (BICS, TRILT, ADL, PDL) we observed no abnormally-elevated EMG activity anywhere in the workspace, consistent with static stretch reflex threshold normally established beyond (i.e., greater than) the muscle’s longest length over the joint’s passive range of motion. In other muscles (TRILG, PECS) we observed substantially-elevated EMG activity throughout the entire workspace, consistent with a static stretch reflex threshold abnormally established much shorter than the muscle’s shortest physiological length. Finally, we observed some muscles (BRD, BICL) that exhibited abnormally elevated responses to passive limb displacement in a way that was sensitive to limb configuration such that activation was greater when the muscle was held at longer lengths than when held at shorter lengths (Fig 8). These results are consistent with the abnormal setting of static stretch reflex thresholds within the joint’s normal range of motion. Important questions for future study are whether these passive bias phenomena may compromise volitional control, and if so, whether it may be possible to mitigate their effects, for example, through training to counter the bias forces using strategic antagonist co-contraction.

### Limitations and future directions

The current study has several limitations. One is the amount of time required to obtain the postural bias maps using the procedure described in the current study. As we have shown however, measured forces and muscle activities achieve steady state 2 seconds after the end of passive displacement of the hand. We also found very robust results when we transported the arm at 90°/s. Consequently, we suggest that robust posture maps could be obtained with brief holding periods between passive displacements performed at this fast speed, thereby reducing testing time by at least one order of magnitude. Another limitation is that we only assessed postural bias in the horizontal plane with the hand supported against gravity by the robot. Although a full 3D volumetric assessment would be intriguing to examine, obtaining it would undoubtedly take a long time, even with abbreviated holding times and relatively fast transport speeds.

Another limitation pertains to the fact that we constructed our postural bias maps only considering passive responses. The postural biases identified here may be very different from those that would be mapped out if the limb were active under volitional control. One possible way to determine if this were so would be to require subjects in a future study to generate low levels of cued co-contraction 2 or 3 seconds after the passively-displaced limb comes to rest at each desired location throughout the workspace. (By co-contraction, we mean here the condition where the subject would generate a minimum of say, 5% MVIC activity in selected flexor/extensor pairs spanning the joint of interest such as the elbow.) Although the resulting hand forces and muscle activities would be modulated by factors including the imbalance of weakness across flexor and extensor muscles and deficits in the coordination of the muscle pairs, comparison of the postural bias maps obtained after passive displacement and during active generation of modest coactivations could yield insight into whether volitional control is constrained in a manner consistent with an impact of postural biases.

## Acknowledgements

This work was supported in part by grants from the National Institutes of Health NICHD R01HD053727 and R15HD093086. The authors thank Drs. John Krakauer and Alkis Hadjiosif for insightful comments on a prior version of this work, and Dr. Leigh A. Mrotek and Ms. Ella Pomplun for valuable assistance in manuscript preparation.

